# Homeostatic and tumourigenic activity of SOX2+ pituitary stem cells is controlled by the LATS/YAP/TAZ cascade

**DOI:** 10.1101/492165

**Authors:** Emily J. Lodge, Alice Santambrogio, John P. Russell, Paraskevi Xekouki, Thomas S. Jacques, Randy L. Johnson, Selvam Thavaraj, Stefan R. Bornstein, Cynthia L. Andoniadou

**Affiliations:** Centre for Craniofacial and Regenerative Biology, Faculty of Dentistry, Oral & Craniofacial Sciences, King’s College London, Floor 27 Tower Wing, Guy’s Campus, London, SE1 9RT, United Kingdom; Division of Diabetes & Nutritional Sciences, Faculty of Life Sciences & Medicine, King’s College London, London SE1 1UL, United Kingdom; Department of Medicine III, University Hospital Carl Gustav Carus, Technische Universität Dresden, Dresden 01307, Germany; Department of Endocrinology, King’s College Hospital NHS Foundation Trust, London, SE5 9RS, UK; UCL GOS Institute of Child Health and Great Ormond Street Hospital for Children NHS Foundation Trust, London, WC1N 1EH, UK; Department of Cancer Biology, The University of Texas, MD Anderson Cancer Center, 1515 Holcombe Blvd., Houston, Texas 77030, USA; Centre for Oral, Clinical and Translational Sciences, Faculty of Dentistry, Oral & Craniofacial Sciences, King’s College London, London SE1 9RT, UK

**Keywords:** pituitary stem cell, SOX2, Hippo, YAP, pituitary tumour

## Abstract

SOX2 positive pituitary stem cells (PSCs) are specified embryonically and persist throughout life, giving rise to all pituitary endocrine lineages. We have previously shown the activation of the MST/LATS/YAP/TAZ signalling cascade in the developing and postnatal mammalian pituitary. Here, we investigate the function of this pathway during pituitary development and in the regulation of the SOX2 cell compartment. Through loss- and gain-of-function genetic approaches, we reveal that restricting YAP/TAZ activation during development is essential for normal organ size and specification from SOX2+ PSCs. Postnatal deletion of LATS kinases and subsequent upregulation of YAP/TAZ leads to uncontrolled clonal expansion of the SOX2+ PSCs and disruption of their differentiation, causing the formation of non-secreting, aggressive pituitary tumours. In contrast, sustained expression of YAP alone results in expansion of SOX2+ PSCs capable of differentiation and devoid of tumourigenic potential. Our findings identify the LATS/YAP/TAZ signalling cascade as an essential component of PSC regulation in normal pituitary physiology and tumourigenesis.

## INTRODUCTION

SOX2 is crucial transcription factor involved in the specification and maintenance of multiple stem cell populations in mammals. Pituitary stem cells express SOX2 and contribute to the generation of new endocrine cells during embryonic development and throughout postnatal life ^1, 2^. The pituitary gland is composed of three parts, the anterior, intermediate and posterior lobes (AL, IL and PL, respectively). The AL and IL contain hormone-secreting cells, which are derived from an evagination of the oral ectoderm expressing SOX2, termed Rathke’s pouch (RP). SOX2+ cells, both in the embryonic and adult pituitary, can differentiate into three endocrine cell lineages, which are marked by transcription factors PIT1 (POU1F1)^3^, TPIT (TBX19)^4^ and SF1 (NR5A1)^5^, and differentiate into hormone-secreting cells (somatotrophs, lactotrophs, thyrotrophs, corticotrophs, melanotrophs and gonadotrophs, which express growth hormone, prolactin, thyrotropin, adrenocorticotropin, gonadotropin and melanotropin, respectively). SOX2+ PSCs acquire PROP1 and SOX9 expression embryonically^2, 6^ and are become highly proliferative during the first 2-3 weeks of life, in concordance with major organ growth, after which they reach a steady low proliferative capacity that contributes to maintain normal homeostasis and physiological adaptation of the pituitary gland ^7, 8^.

Contrary to other organs, where somatic stem cells are shown to be able to become transformed into cancer stem cells, the roles of SOX2+ PSCs in tumourigenesis remain poorly understood, possibly due to the patchy knowledge of the pathways regulating SOX2+ PSC fate and proliferation. Pituitary tumours are common in the population, representing 10-15% of all intracranial neoplasms ^9, 10^. Adenomas are the most common adult pituitary tumours, classified into functioning, when they secrete one or more of the pituitary hormones, or non-functioning if they do not secrete hormones. In children, adamantinomatous craniopharyngioma (ACP) is the most common pituitary tumour. Targeting oncogenic beta-catenin in SOX2+ PSCs in the mouse generates clusters of senescent SOX2+ cells that induce tumours resembling ACP in a paracrine manner, i.e. the tumours do not derive from the targeted SOX2+ PSCs ^1, 11^. Up to 15% of adenomas and 50% of ACP display aggressive behaviour with invasion of nearby structures including the hypothalamus and visual tracts, associated with significant morbidity and mortality ^12^. Pituitary carcinomas exhibiting metastasis are rare but can develop from benign tumours ^13, 14, 15^. Whether SOX2+ cells can cell autonomously contribute to pituitary neoplasia has not been hitherto demonstrated.

The Hippo pathway controls stem cell proliferation and tumourigenesis in several organs such as in the liver^16, 17^, intestines^18^ and lung^19, 20^. In the core phosphorylation cascade, MST1/2 kinases phosphorylate and activate LATS1/2 serine/threonine-protein kinases, which in turn phosphorylate co-activators Yes-associated protein (YAP) and WW domain-containing transcription regulator protein 1 (WWTR1, a.k.a. TAZ) that are subsequently inactivated through degradation and cytoplasmic retention ^21^. Active YAP/TAZ associate with TEAD transcription factors, promoting the transcription of target genes such as *Cyr61* and *Ctgf* ^22, 23, 24^. YAP/TAZ have been shown to promote proliferation and the stem cell state in several organs, and can also lead to transformation and tumour initiation when overexpressed^25, 26, 27^. The involvement of YAP/TAZ in the function of tissue-specific SOX2+ stem cells during development and homeostasis has not been shown. We previously reported strong nuclear localisation of YAP and TAZ exclusively in SOX2+ stem cells of developing Rathke’s pouch and the postnatal anterior pituitary of mice and humans, and enhanced expression in human pituitary tumours composed of uncommitted cells, including ACPs and null-cell adenomas^28, 29^. In these populations we detected phosphorylation of YAP at serine 127 (S127) indicating LATS kinase activity. Together these point to a possible function for LATS/YAP/TAZ in normal pituitary stem cells and during tumourigenesis. Here, we have combined genetic and molecular approaches to reveal that deregulation of the pathway can promote and maintain the SOX2+ PSC fate under physiological conditions and that major disruption of this axis transforms SOX2+ PSCs into cancer-initiating cells giving rise to aggressive tumours.

## RESULTS

### Sustained conditional expression of YAP during development promotes SOX2+ PSC fate

To determine if YAP and TAZ function during embryonic development of the pituitary, we used genetic approaches to perform gain- and loss-of-function experiments. We first expressed a constitutive active form of YAP(S127A) using the *Hesx1-Cre* driver, which drives *Cre* expression in Rathke’s pouch (RP) and the hypothalamus primordium from 9.5dpc, regulated by administration of doxycycline through the reverse tetracycline-dependent transactivator (rtTA) system (*R26^rtTA/+^*; see Methods for details, Scheme Fig1A). Analyses were restricted to embryonic time points. As expected, we confirmed accumulation of total YAP protein but not of TAZ or pYAP(S127), throughout the developing pituitary and hypothalamus of *Hesx1^Cre/+^;R26^rtTA/+^;Col1a1^tetO-Yap^1^/+^* embryos at 15.5dpc, but not of *Cre*- negative controls (Fig1B, SFig1A). Likewise, the YAP downstream target *Cyr61* (Fig1B) was also upregulated. Morphologically, YAP-TetO mutants displayed a dysplastic anterior pituitary, which was more medially compacted and lacked a central lumen, making it difficult to distinguish between the developing anterior and intermediate lobes (Fig 1C).

**Figure 1.**
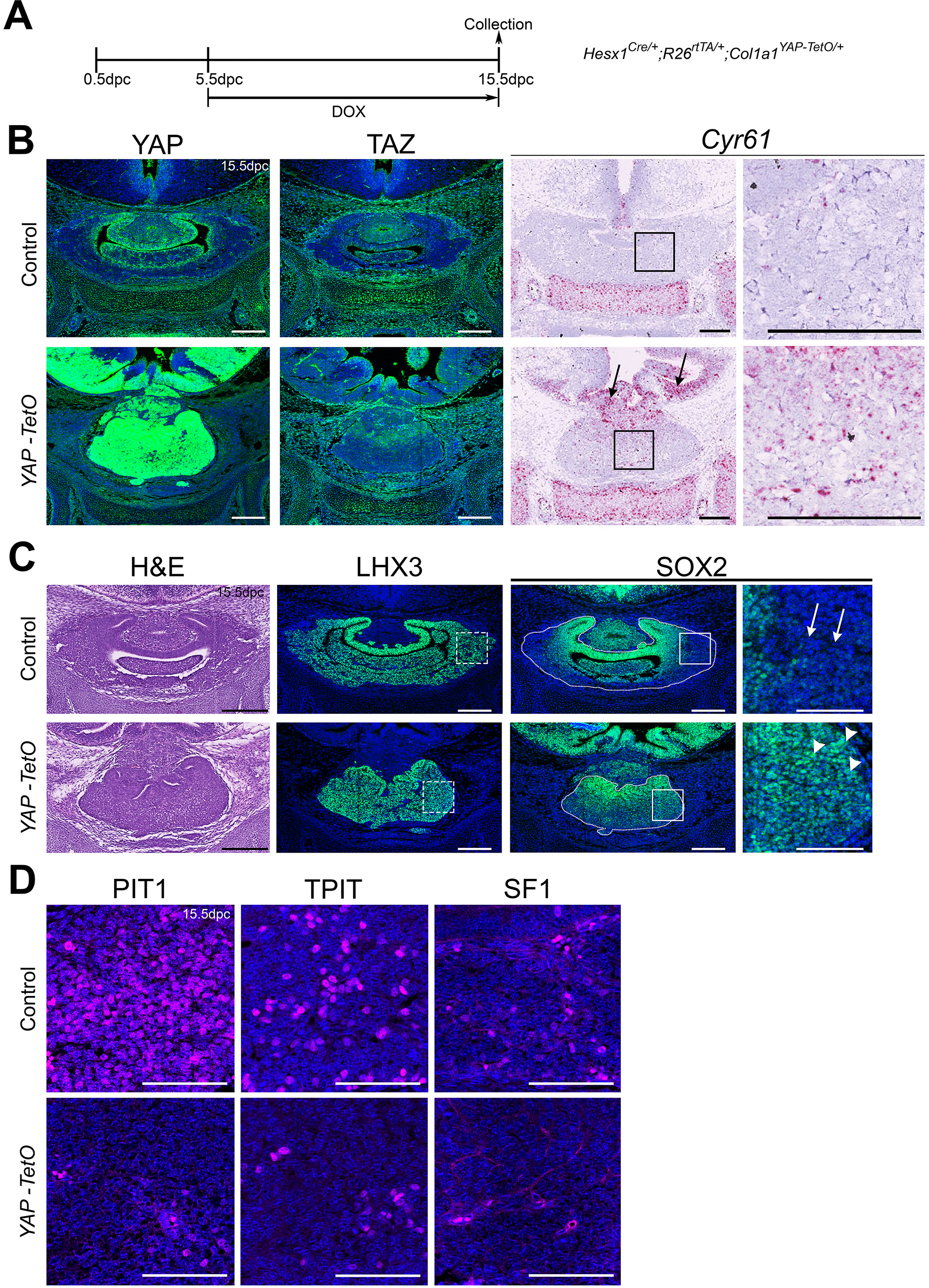
Regulation of YAP is required for normal morphogenesis and lineage commitment during pituitary development. **A.** Schematic outlining the time course of doxycycline (DOX) treatment administered to pregnant dams from *Hesx1^Cre/+^* x *R26^rtTA/rtTA^;Col1a1^tetO-Yap1^/tetO-Yap^1^* crosses for the embryonic induction of YAP(S127A) expression in *Hesx1^Cre/+^;R26^rtTA/+^;Col1a1^tetO-Yap^1^/+^* (*YAP-TetO*) mutant embryos as well as controls that do not express YAP(S127A) (*Hesx1^+/+^;R26^rtTA/+^;Col1a1^tetO-Yap^1^/+^* controls shown here). **B.** Immunofluorescence staining against YAP and TAZ on frontal pituitary sections at 15.5dpc confirms accumulation of YAP protein in *YAP-TetO* compared to control sections, but no increase in TAZ levels. RNAscope mRNA *in situ* hybridisation against the YAP/TAZ target *Cyr61* confirms an increase in transcripts in the anterior pituitary as well as the hypothalamus where the Cre is also active (arrows). **C.** Haematoxylin and eosin staining of frontal pituitary sections from 15.5dpc control and *YAP-TetO* embryos showing pituitary dysmorphology in mutants. Immunofluorescence staining for LHX3 to mark anterior pituitary tissue and SOX2 to mark pituitary progenitors shows the persistence of SOX2 protein in lateral regions of the gland in *YAP-TetO* mutants (arrowheads) when they have lost SOX2 expression in controls (arrows) (magnified boxed region in SOX2, corresponding to dashed box in LHX3). **D.** Immunofluorescence staining for lineage-committed progenitor markers PIT1, TPIT and SF1 reveals very few cells expressing commitment markers in *YAP-TetO* compared to control. Scale bars 100µm, 50µm in magnified boxed regions in C. See Also Supplementary Figure 1.

Immunofluorescence staining against SOX2 at 15.5dpc demonstrated loss of SOX2 in the most lateral regions of control pituitaries (arrows in Fig. 1C), where cells are undergoing commitment; yet mutant pituitaries had abundant SOX2 positive cells in the most lateral regions (arrowheads in Fig1C). Immunostaining for LHX3, which is expressed in the developing anterior pituitary ^30^, was used to demarcate AL and IL tissue. Staining using antibodies against lineage markers PIT1, TPIT and SF1 revealed a concomitant reduction in committed cell lineages throughout the gland (Fig 1D; PIT1 0.35% in mutants compared to 30.21% in controls (Student’s t-test *P*<0.0001, n=3 for each genotype), TPIT 1.03% in mutants compared to 9.81% in controls (Student’s t-test *P*=0.0012, n=3 for each genotype), SF1 0.34% in mutants compared to 4.14% in controls (Student’s t-test *P*=0.0021, n=3 for each genotype)). We therefore conclude that sustained activation of YAP prevents lineage commitment and is sufficient to maintain the progenitor state during embryonic development.

Next, we generated embryos null for TAZ and conditionally lacking YAP in the *Hesx1* expression domain (SFig1B-E). *Hesx1^Cre/+^;Yap^fl/fl^;Taz^-l-^* double mutants were obtained at expected ratios during embryonic stages until 15.5dpc, however the majority of *Taz^-/-^* mutants with or without compound *Yap* deletions showed lethality at later embryonic and early postnatal stages ^31^ (Table 1). The developing pituitary gland of *Hesx1^Cre/+^;Yap^fl/fl^;Taz^-l-^* double mutants appeared largely normal at 13.5dpc by histology (SFig1B). Immunostaining against SOX2 to mark embryonic progenitors and postnatal stem cells did not reveal differences in the spatial distribution of SOX2+ cells between double mutants compared to controls (*Hesx1^+/+^;Yap^fl/fl^;Taz^+/+^* and *Hesx1^+/+^;Yap^fl/fl^;Taz^+/-^*) at 13.5dpc, 16.0dpc (SFig1C) or P28, even in regions devoid of both TAZ and active YAP (SFig1D). This suggests that YAP/TAZ are not required for SOX2+ cell specification or survival. Likewise, analysis of commitment markers PIT1and SF1 as well as ACTH to mark the TPIT lineage, did not show any differences between genotypes (SFig1E). Together, these data suggest there is no critical requirement for YAP and TAZ during development for the specification of SOX2+ cells or lineage commitment, but that YAP functions to promote the SOX2 cell identity.

### LATS, but not MST, kinases are required for normal pituitary development and differentiation

Since sustained activation of YAP led to an embryonic phenotype, we reasoned that YAP/TAZ need to be regulated during embryonic development. To determine if MST and LATS kinases are important in YAP/TAZ regulation we carried out genetic deletions in the pituitary.

Conditional deletion of *Mst1* and *Mst2* (official names, *Stk4* and *Stk3*) in *Hesx1^Cre/+^;Mst1^fl/fl^;Mst2^fl/fl^* embryos did not lead to a pituitary phenotype (SFig2). Mutant pituitaries were macroscopically normal at birth (SFig2A), and showed comparable expression patterns of TAZ, YAP, pYAP to controls lacking *Cre*, without distinct accumulation of YAP or TAZ (SFig2B). The distribution of SOX2+ cells was comparable between mutants and controls (SFig2B). Normal lineage commitment was evident by immunofluorescence staining for PIT1, TPIT and SF1 at P10 (SFig2C). Mutant animals remained healthy and fertile until P70, at which point pituitaries appeared histologically normal (SFig2D). Since deletion of *Mst1/2* at embryonic stages does not affect embryonic or postnatal pituitary development, we conclude these kinases are not critical for YAP/TAZ regulation in the pituitary.

We next focused on perturbing LATS kinase function, as we have previously shown strong expression of *Lats1* in the developing pituitary and postnatal kinase activity in SOX2+ stem cells^28^. However, *Hesx1^Cre/+^;Lats1^fl/fl^* embryos showed unaffected pituitary development and normal localisation and levels of YAP and TAZ as assessed by immunofluorescence (SFig3A,B) when compared with controls. mRNA *in situ* hybridisation against *Lats2* at P2, revealed abundant *Lats2* transcripts upon conditional deletion of *Lats1*, suggesting a compensatory upregulation of *Lats2* in the absence of LATS1 (SFig3C), similar to previous reports of elevated YAP/TAZ signalling inducing *Lats2* expression^32^.

To overcome potential functional redundancy, we deleted both *Lats1* and *Lats2* in RP. Deletion of *Lats2* alone (*Hesx1^Cre/+^;Lats2^fl/fl^*), did not reveal any developmental morphological anomalies (SFig3D) and pups were identified at normal Mendelian proportions (Table 2). Similarly, deletion of any three out of four *Lats* alleles did not affect pituitary development and were identified at normal ratios, similar to other tissues ^33^. Homozygous *Hesx1^Cre/+^;Lats1^fl/fl^;Lats2^fl/fl^* mutants were identified at embryonic stages at reduced Mendelian ratios and were absent at P0-P2, suggesting embryonic and perinatal lethality (Table 2).

Haematoxylin/eosin staining of the developing pituitary gland in *Hesx1^Cre/+^;Lats1^fl/fl^;Lats2^fl/fl^* mutants revealed overgrowth of RP by 13.5dpc compared to controls lacking *Cre* (Fig2A, n=4). Total TAZ and YAP proteins accumulated throughout the developing gland in double mutants (arrowheads) but only in the SOX2+ periluminal epithelium of controls (arrows). The same regions showed a marked reduction in pYAP-S127 staining, which is observed in SOX2+ cells of the control (Fig2A). These findings are in line with LATS1/2 normally regulating YAP and TAZ in the pituitary and demonstrate successful deletion in RP. The mutant pituitary was highly proliferative (Fig2B; Ki-67 index average 26.98% ±6.0 SD in control versus 80.14% ±4.09 SD in the double mutant, *P*≤0.0001, Student’s *t*-test) and the majority of cells expressed SOX2 (Fig2A,C) but not SOX9 (Fig2B).

**Figure 2.**
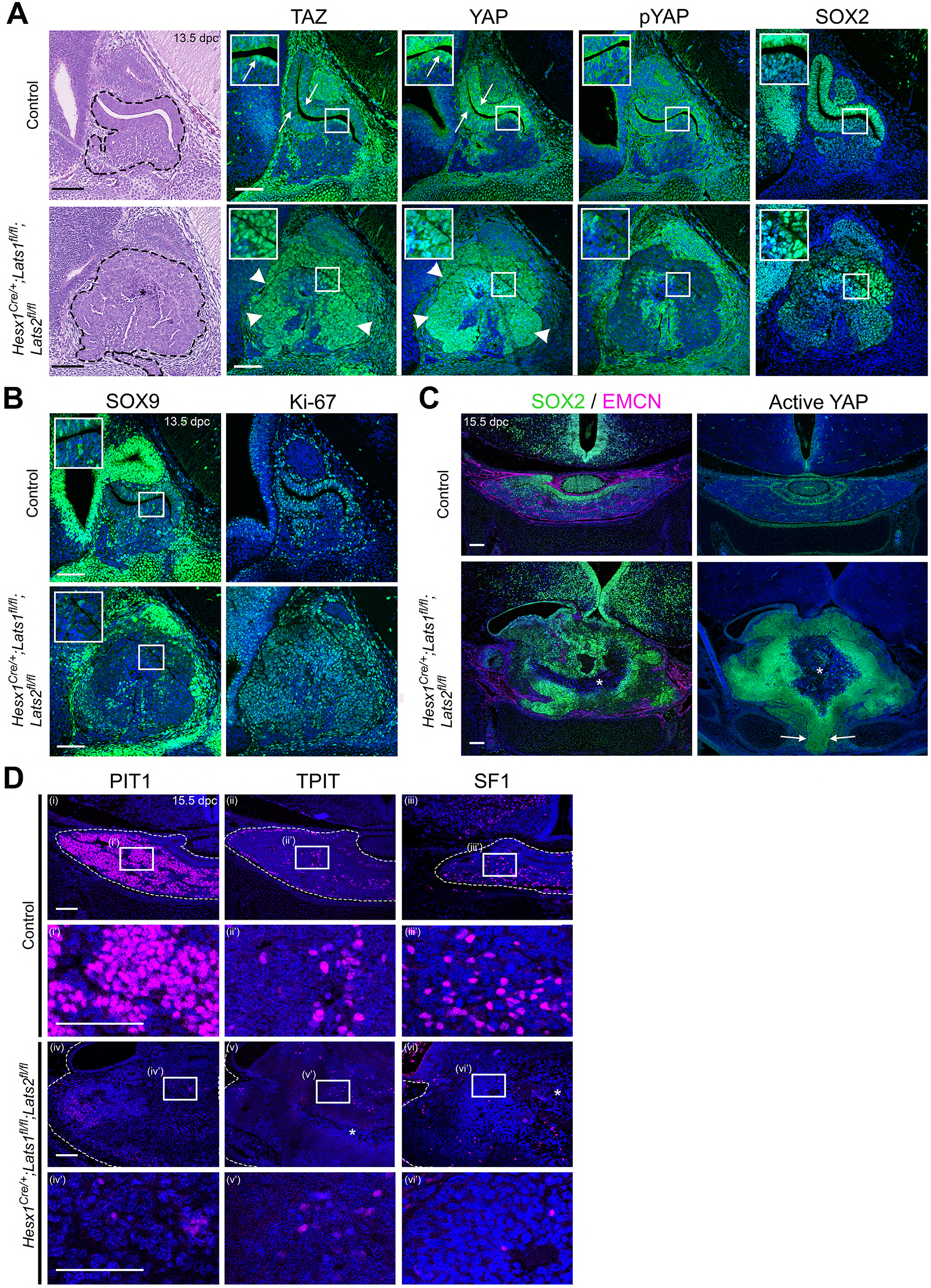
Pituitary-specific deletion of *Lats1* and *Lats2* during development leads to pituitary overgrowth and defects in lineage commitment. **A.** Haematoxylin and eosin staining on sagittal sections from *Hesx1^Cre/+^;Lats1^fl/fl^;Lats2^fl/fl^* (mutant) and *Hesx1^+/+^;Lats1^fl/fl^;Lats2^fl/fl^* (control) embryos at 13.5dpc reveals anterior pituitary dysmorphology and overgrowth in mutants (dashed outline). Immunofluorescence staining for TAZ, YAP and pYAP reveals accumulation of TAZ and YAP in overgrown mutant tissue (arrowheads, normal epithelial expression indicated by arrows in control) and lack of staining for pYAP (S127). Immunofluorescence for SOX2 shows the presence of SOX2+ progenitors throughout the abnormal tissue in mutants. **B.** Immunofluorescence staining for late progenitor marker SOX9 shows localisation in few cells of the pituitary of mutants at 13.5dpc. Immunofluorescence staining for Ki-67 indicates cycling cells throughout the mutant pituitary. **C.** Immunofluorescence staining for SOX2 and Endomucin (EMCN) on frontal pituitary sections at 15.5dpc shows expansion of the SOX2+ progenitor compartment compared to controls and a reduction in vasculature marked by endomucin. Immunofluorescence for non-phosphorylated (Active) YAP shows strong expression throughout the mutant gland compared to the control. Areas of necrosis in mutant tissue indicated by asterisks. Ventral overgrowth extending into the oral cavity between the condensing sphenoid bone indicated by arrows. **D.** Immunofluorescence staining for lineage-committed progenitor markers PIT1, TPIT and SF1 reveals only sporadic cells expressing commitment markers in *Hesx1^Cre/+^;Lats1^fl/fl^;Lats2^fl/fl^* mutants compared to controls. Boxes showing magnified regions. Dashed lines demarcate anterior pituitary tissue. Scale bars 100µm.

By 15.5dpc the pituitary was grossly enlarged and exerting a mass effect on the brain, had cysts and displayed areas of necrosis (asterisks Fig2, SFig3E n=5). Staining for endomucin to mark blood vessels revealed poor vascularisation in *Hesx1^Cre/+^;Lats1^fl/fl^;Lats2^fl/fl^* mutants compared to the ample capillaries seen in the control (Fig2C), which may account for the necrosis. We frequently observed ectopic residual pituitary tissue at more caudal levels, reaching the oral epithelium and likely interfering with appropriate fusion of the sphenoid, similar to other pituitary hyperplasia phenotypes (arrow Fig2C) ^34, 35, 36^. Immunofluorescence to detect active (non-phosphorylated) YAP revealed abundant staining throughout the pituitary at 15.5dpc, compared to the control where active YAP localises in the SOX2 epithelium (Fig2C). Immunofluorescence using specific antibodies against lineage commitment markers PIT1, TPIT and SF1 at 15.5dpc revealed very few cells expressing PIT1, TPIT and SF1 in the double mutant (up to ∼ 9%, 4% and 2% respectively, n=3 mutants) compared to controls (∼ 50%, 11% and 6%, respectively in controls, n=5) suggesting failure to commit into the three lineages (Fig2D). These data suggest that the LATS/YAP/TAZ axis is required for normal embryonic development of the anterior pituitary and that LATS1/2 kinases control proliferation of SOX2+ progenitors and their progression into the three committed lineages.

### Loss of LATS kinases results in carcinoma-like murine tumours

Postnatal analysis of *Hesx1^Cre/+^;Lats1^fl/fl^* pituitaries revealed that by P56, despite developing normally during the embryonic period, all glands examined exhibited lesions of abnormal morphology consisting of overgrowths, densely packed nuclei and loss of normal acinar architecture (n=15). To minimise the likely redundancy by LATS2 seen at embryonic stages, we generated *Lats1* mutants additionally haploinsufficient for *Lats2* (*Hesx1^Cre/+^;Lats1^fl/fl^;Lats2^fl/+^*). These pituitaries also developed identifiable lesions accumulating YAP and TAZ (SFig4A), which were observed at earlier time points (P21 n=4), the earliest being 10 days indicating increased severity. The number of lesions observed per animal was similar between the two models at P56 (3-8 per animal). Deletion of *Lats2* alone (*Hesx1^Cre/+^;Lats2^fl/fl^*), which is barely expressed in the wild type pituitary, did not result in any defects (SFig4B). We focused on the *Hesx1^Cre/+^;Lats1^fl/fl^;Lats2^fl/+^* double mutants for further analyses.

Histological examination of *Hesx1^Cre/+^;Lats1^fl/fl^;Lats2^fl/+^* pituitaries confirmed the abnormal lesions were tumours, characterised by frequent mitoses, focal necrosis, and a focal squamous differentiation, as well as the occasional presence of cysts (Fig3A). These lesions were identical to those in *Hesx1^Cre/+^;Lats1^fl/fl^* pituitaries (not shown). These tumours accumulated YAP/TAZ and upregulated expression of targets *Cyr61* and *Ctgf* (Fig5C), confirming the validity of the genetic manipulation (Fig3B). Tumours were also frequently observed in the anterior and intermediate lobe (SFig4C). Analysis of proliferation by Ki-67 immunostaining revealed an elevated mitotic index of 7-28% in tumours (mean 15.46, SEM ±2.74), compared to 2.97% (SEM ±1.2) mean in control pituitaries not carrying the *Lats1* deletion (Fig3C). In keeping with the morphological evidence of epithelial differentiation, the tumours were positive for cytokeratins using AE1/AE3 (multiple keratin cocktail) (SFig4E). Furthermore, the tumours showed focal morphological evidence of squamous differentiation and this was confirmed by p63 staining (SFig4E). In contrast, the tumours did not show immunohistochemical evidence of adenomas and were negative for typical adenoma markers: the neuroendocrine marker synaptophysin and neuron-specific enolase (SFig4F). The lesions were also negative for chromogranin A, a neuroendocrine granule marker often expressed in clinically non-functioning pituitary adenomas, as well as negative for vimentin, characteristic of spindle cell oncocytoma (SFig4F). Moreover, immunostaining against PIT1, TPIT and SF1 showed only sparse positive cells within the lesions, suggesting lack of commitment into endocrine precursors and supporting the undifferentiated nature of the tumour cells (Fig3D). Consistent with a tumourigenic phenotype, and role for LATS1 genomic stabilisation ^37^, staining for gamma-H2A.X detected elevated DNA damage in cells of the mutant pituitaries compared with controls (SFig4D). The absence of adenoma or oncocytoma markers together with the histological appearance, observation of focal necrosis and a high mitotic index support the features of squamous carcinoma.

**Figure 3.**
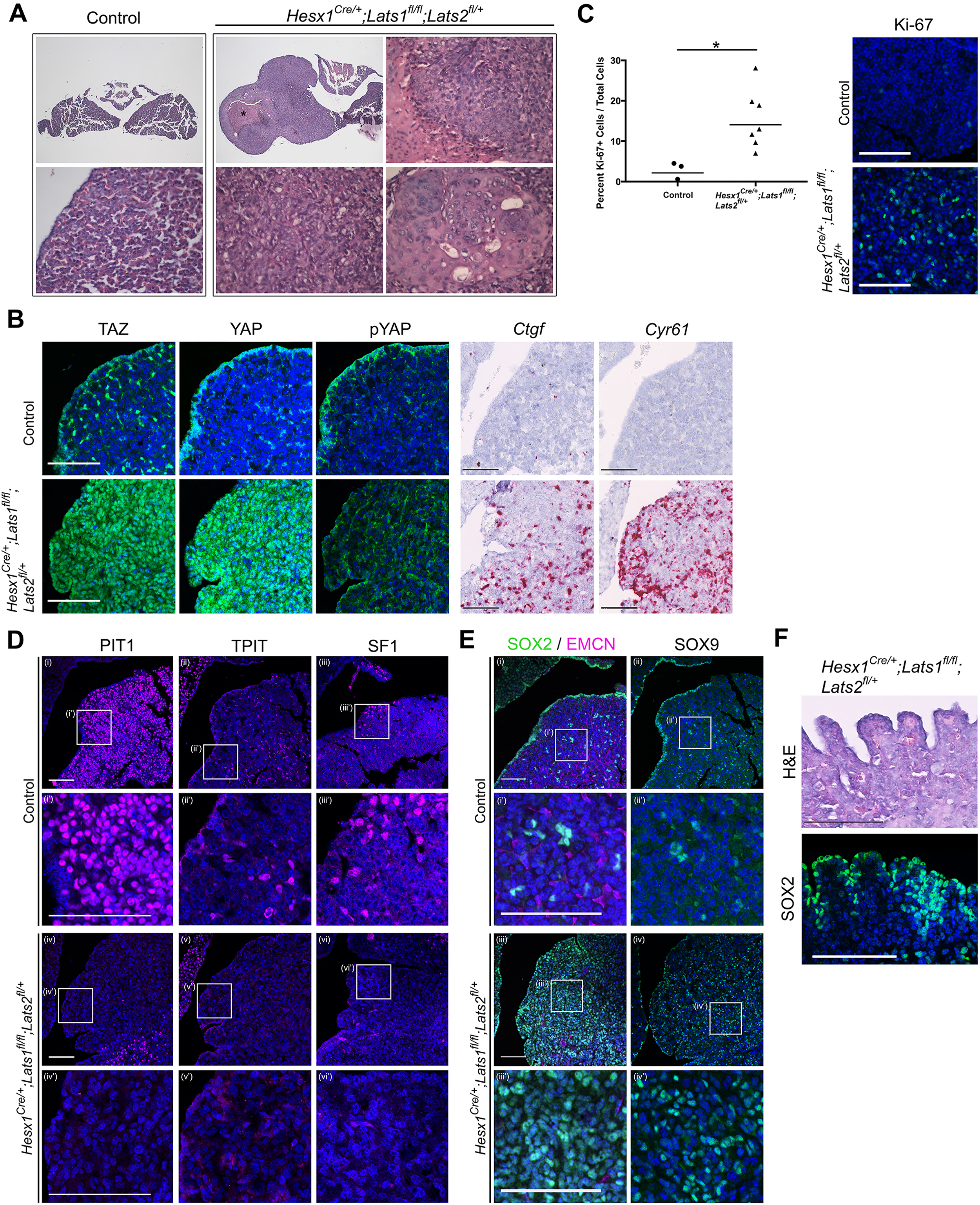
Pituitary specific loss of *Lats1* leads to tumour formation. **A.** Haematoxylin and eosin staining of frontal sections from *Hesx1^Cre/+^;Lats1^fl/fl^;Lats2^fl/+^* (mutant) and control pituitaries at P56 demonstrates overgrown tumourigenic regions in mutants. These show focal necrosis, cysts and a squamous morphology (magnified regions) not seen in controls. Asterisk indicates necrosis. **B.** Immunofluorescence staining for TAZ, YAP and pYAP(S127) show accumulation of TAZ and YAP but not pYAP in the mutant but not in the control. RNAscope mRNA *in situ* hybridisation against YAP/TAZ targets *Ctgf* and *Cyr61* reveals an increase in transcripts on mutant tissue compared to control. **C.** Graph of the proliferation index in control and mutant samples at P56 shows a significant increase in cycling cells in the *Hesx1^Cre/+^;Lats1^fl/fl^;Lats2^fl/+^* mutant pituitaries compared to controls (control percentage Ki-67: 2.967±1.2 SEM, n=3; mutant: 15.46±2.74 n=7. P value: 0.0217 (*), two-tailed t test). Images show representative examples of Ki-67 immunofluorescence staining. **D.** Immunofluorescence staining for lineage-committed progenitor markers PIT1, TPIT and SF1 shows the near absence of committed cells in tumours. **E.** Immunofluorescence staining for pituitary stem cell markers SOX2 and SOX9 reveal that tumour lesions have abundant positive cells compared to the control, whilst endomucin (EMCN) staining shows poor vascularisation. **F.** The marginal zone epithelium of *Hesx1^Cre/+^;Lats1^fl/fl^;Lats2^fl/+^* mutant pituitaries develops invaginations as seen by haematoxylin and eosin staining. Immunofluorescence staining against SOX2 shows the maintenance of a single-layered epithelium. Scale bars 100 µm. Boxes indicate magnified regions.

### SOX2 +ve cells are the cell of origin of the tumours

Tumour regions were mostly composed of SOX2 positive cells, a sub-population of which also expressed SOX9 (Fig3E, SFig4A; 85-97% of cells, 7 tumours across 4 pituitaries). Close examination of the marginal zone epithelium, a major SOX2+ stem cell niche of the pituitary, revealed a frequent ‘ruffling’ resembling crypts, likely generated through over-proliferation of the epithelial stem cell compartment (Fig3F). To determine if the cell of origin of the tumourigenic lesions is a deregulated SOX2+ stem cell, we carried our specific deletion of LATS1/2 in postnatal SOX2+ cells using the tamoxifen-inducible *Sox2-CreERT2* driver, combined with conditional expression of membrane-GFP in targeted cells (*Sox2^CreERT2/+^;Lats1^fl/fl^;Lats2^fl/+^;R26^mTmG/+^*).

Tamoxifen induction at P5 or P21, led to abnormal lesions in the anterior pituitary within two months in all cases. We focused our analyses on inductions performed at P5, from which time point all animals developed lesions by P35 (Fig4A). Similar to observations in *Hesx1^Cre/+^;Lats1^fl/fl^;Lats2^fl/+^* animals, these areas strongly accumulated YAP and TAZ (Fig4B), activated expression of targets *Cyr61* and *Ctgf*, displayed ruffling of the AL epithelium (Fig4C) and lacked lineage commitment markers (Fig4D). These lesions showed a similar marker profile to *Hesx1-Cre*-targeted tumours, with positive p63 and AE1/AE3 staining (SFig5A). Lineage tracing confirmed expression of membrane GFP in tumourigenic lesions, characterised by the accumulation of YAP and expansion of SOX2+ cells, suggesting they were solely derived from SOX2+ cells (Fig4E, SFig5B). Taken together, our data support that LATS kinase activity is required to regulate the pituitary stem cell compartment. Loss of LATS1 is sufficient to drive deregulation of SOX2+ pituitary stem cells, generating highly proliferative non-functioning tumours with features of carcinomas.

**Figure 4.**
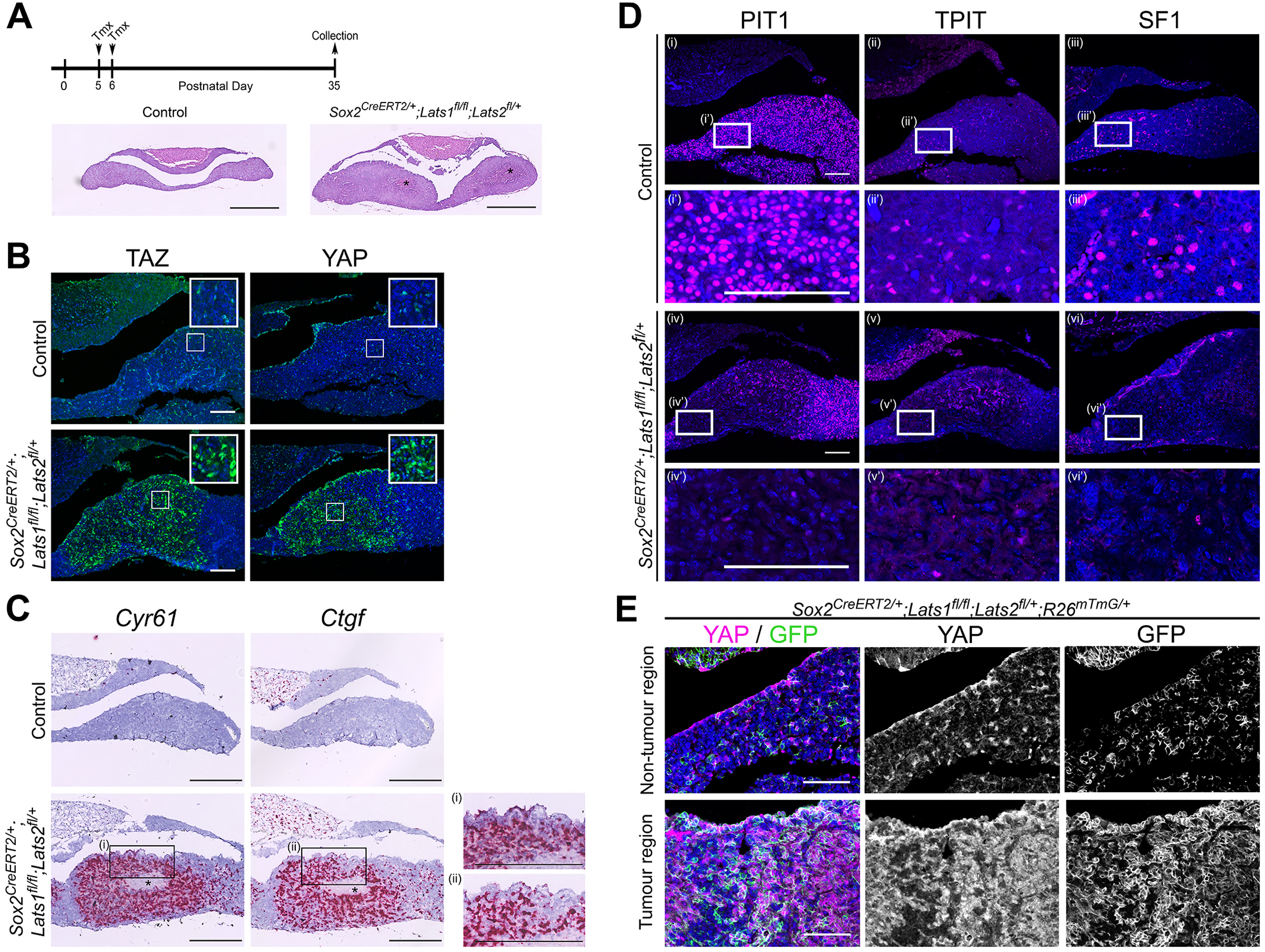
SOX2+ pituitary stem cells are the cell-of-origin of tumours generated in the absence of *Lats1*. **A.** Schematic outlining the experimental time line of inductions in *Sox2^CreERT^2^/+^;Lats1^fl/fl^;Lats2^fl/+^* (mutant) and *Sox2^+/+^;Lats1^fl/fl^;Lats2^fl/+^* (control) animals. Representative images of haematoxylin and eosin staining of frontal sections of control and mutant pituitaries at P35, revealing a hyperplastic anterior pituitary in the mutant with areas of necrosis (asterisks). **B.** Immunofluorescence staining reveals tumourigenic lesions in *Sox2^CreERT^2^/+^;Lats1^fl/fl^;Lats2^fl/+^* that display increased levels of TAZ and YAP staining compared to the control. **C.** RNAscope mRNA *in situ* hybridisation against *Ctgf* and *Cyr61* shows elevated transcripts in tumourigenic lesions. Insets (i) and (ii) show invaginations in the epithelium of the mutant. **D.** Immunofluorescence staining for lineage-committed progenitor markers PIT1, TPIT and SF1 showing a reduction in staining in tumourigenic lesions compared to control pituitaries. **E.** Lineage tracing of SOX2+ cells in *Sox2^CreERT^2^/+^;Lats1^fl/fl^;Lats2^fl/+^R26^mTmG/+^* reveals that tumour regions accumulating YAP as seen by immunofluorescence, are composed of GFP+ cells at P35. Scale bars 500µm in A; 100µm in B, D, E; 250µm in C.

### YAP expression is sufficient to activate pituitary stem cells

Conditional deletion of LATS1/2 kinases in the pituitary has revealed how these promote an expansion of SOX2+ve stem cells in the embryonic and postnatal gland at the expense of differentiation. To establish if this effect was mediated through YAP alone, we used the tetracycline-controlled conditional YAP-TetO system to promote YAP (S127A) protein levels in postnatal pituitaries of *Hesx1^Cre/+^;R26^rtTA/+^;Col1a1^tetO-Yap^1^/+^* mice. We treated YAP-TetO animals with doxycycline from P21 to P105 (12 week treatment, Fig5A). We did not observe the formation of tumours at any stage analysed (n=12, SFig6A). Similarly, we did not observe the formation of lesions when treating from P5. This is in contrast with the unequivocal tumour formation observed in *Sox2^CreERT^2^/+^;Lats1^fl/fl^;Lats2^fl/+^*mice. Elevation of YAP protein levels was confirmed following three weeks of doxycycline treatment (P42), displaying patchy accumulation, likely a result of genetic recombination efficiencies (Fig5B). Consistent with pathway activation, there was robust elevation in the expression of transcriptional targets *Cyr61* and *Ctgf* following treatment (SFig6B) and there was no elevation in phosphorylated inactive YAP (Fig5B).

**Figure 5.**
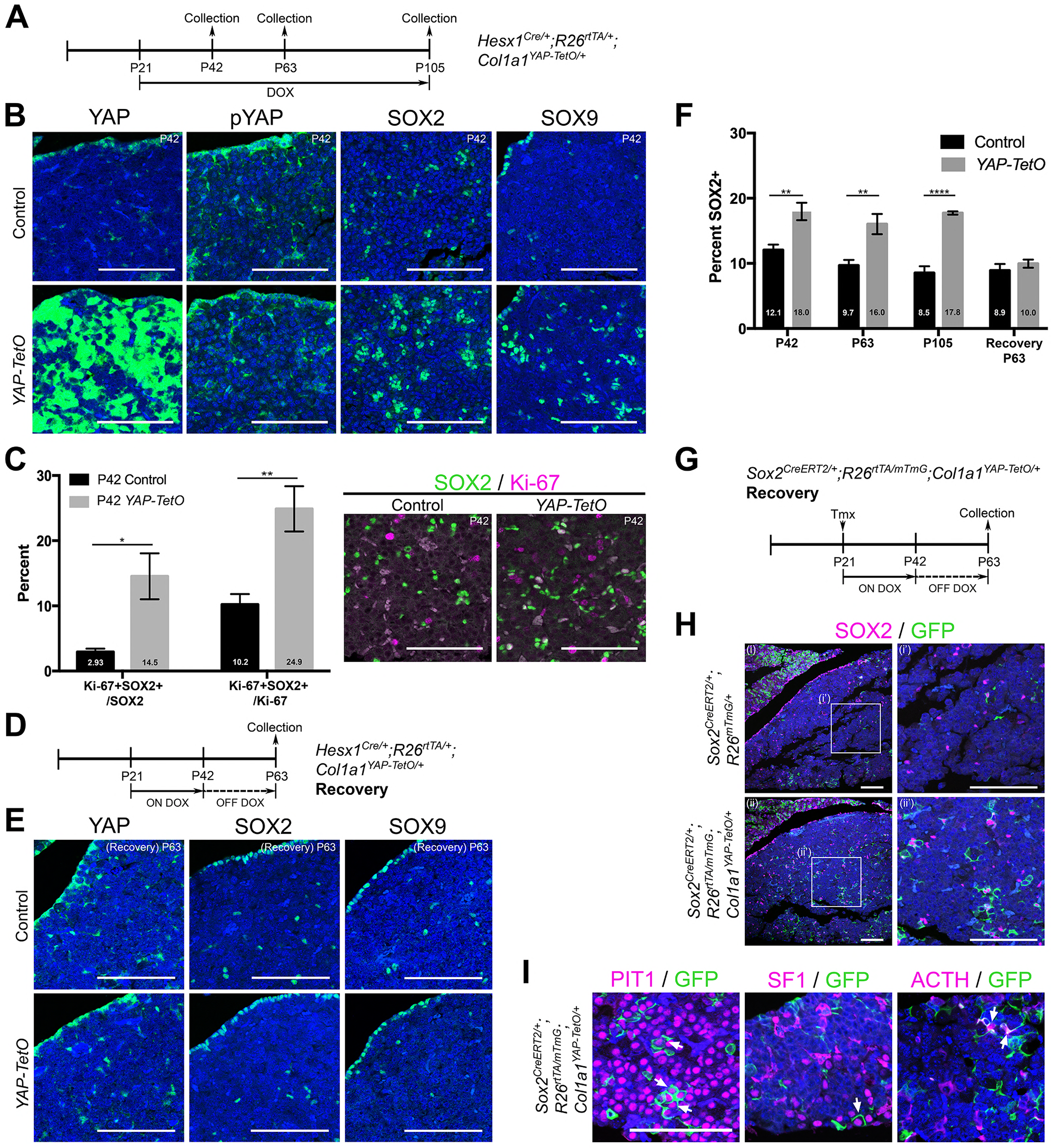
Postnatal expression of constitutively active YAP increases leads to an activation of SOX2+ pituitary stem cells. **A.** Schematic outlining the time course of doxycycline (DOX) treatment administered to *Hesx1^Cre/+^;R26^rtTA/+^;Col1a1^tetO-Yap^1^/+^* (*YAP-TetO*) and *Hesx1^+/+^;R26^rtTA/+^;Col1a1^tetO-Yap^1^/+^* controls to drive expression of YAP-S127A in mutant pituitaries. **B.** At P42 (3 weeks of treatment), immunofluorescence staining on frontal anterior pituitary sections detects strong total YAP expression in *YAP-TetO* mutants compared to the control and no increase in pYAP(S127). Immunofluorescence for SOX2 and SOX9 reveals an expanded population of stem cells in *YAP-TetO* compared to control (quantification in F). **C.** Graph showing the percentage of double Ki-67+SOX2+ cells as a proportion of the total SOX2+ (P=0.027 (*)) or Ki-67+ (P=0.006 (**)) populations at P42 (n=3 pituitaries per genotype). There is an increase in the numbers of cycling SOX2 cells in *YAP-TetO* mutant compared to controls. The image shows a representative example of double immunofluorescence staining against Ki-67 and SOX2 in a control and *YAP-TetO* section. **D.** Schematic outlining the time course of doxycycline (DOX) treatment administered to *Hesx1^Cre/+^;R26^rtTA/+^;Col1a1^tetO-Yap^1^/+^* (*YAP-TetO*) and *Hesx1^+/+^;R26^rtTA/+^;Col1a1^tetO-Yap^1^/+^* controls to drive expression of YAP-S127A in mutant pituitaries for three weeks, followed by a three-week recovery period in the absence of DOX. **E.** Immunofluorescence staining against YAP, SOX2 and SOX9 on control and *YAP-TetO* pituitaries treated as in D, shows comparable expression of YAP, SOX2 and SOX9 between genotypes. **F.** Graph of quantification of SOX2+ cells as a percentage of total nuclei in control and *YAP-TetO* pituitaries at P42 P=0.0014 (**); P63 P=0.0044 (**); P105 P<0.0001(****) (n=3 pituitaries per genotype). Following the Recovery treatment scheme in D, there is no significant difference in the numbers of SOX2+ cells between genotypes. **G.** Schematic outlining the time course of tamoxifen induction and doxycycline (DOX) treatment administered to *Sox2^CreERT^2^/+^;R26^rtTA/mTmG^;Col1a1^tetO-Yap^1^/+^* (mutant) and *Sox2^CreERT2/+^;R26^mTmG/+^;Col1a1^+/+^* (control) animals to drive expression of YAP-S127A in SOX2+ cells of mutants. **H.** Lineage tracing of SOX2+ cells and immunofluorescence staining against SOX2 and GFP shows an expansion of GFP+ cells compared to controls at P63, where a proportion of cells are double-labelled. **I.** Immunofluorescence staining against commitment markers PIT1, SF1 and terminal differentiation marker ACTH (TPIT lineage) together with antibodies against GFP detects double-labelled cells (arrows) across all three lineages in *Sox2^CreERT2/+^;R26^rtTA/mTmG^;Col1a1^tetO-Yap^1^/+^* pituitaries following the recovery period. Scale bars 100µm. Data in C. and F. represented as mean ± SEM, analysed with Two-Way ANOVA with Sidak’s multiple comparisons.

Immunofluorescence against SOX2 demonstrated a significant increase in the number of SOX2+ cells as a proportion of the anterior pituitary (Fig5B,F; 18.0% compared to 12.1% in controls, *P*=0.0014), a finding recapitulated by SOX9 that marks a subset of the SOX2 population (Fig5B). This increase in the percentage of SOX2+ cells was maintained at all stages analysed (Fig5F) and did not affect the overall morphology of the pituitary. At P42 we observed a significant increase in proliferation among the SOX2+ pituitary stem cells from 3% in controls to 15% in mutants (*P*=0.027). SOX2+ cells make up 11% of all cycling cells (Ki-67%) in normal pituitaries, however in mutants this increased to 24%, suggesting a preferential expansion of the SOX2+ population, rather than an overall increase in proliferation (Fig5C). No additional marked differences were observed in samples analysed at P63 (6 weeks of treatment, n=3), however longer treatment (P21 to P105) resulted in sporadic regions of expanded SOX2+ cells (SFig6C). These regions did not express the commitment marker PIT1 and were identifiable by haematoxylin/eosin staining. In contrast to tumour lesions generated following loss of LATS kinases, these were not proliferative, were positive for pYAP and did not accumulate high levels of YAP/TAZ (n=6 lesions). Together these results suggest that the sustained expression of constitutive active YAP can activate the proliferation of SOX2 stem cells, but in contrast to deletion of LATS1, this alone is not oncogenic.

To establish if the expansion of pituitary stem cells following forced expression of YAP is reversible, we administered doxycycline to YAP-TetO animals for three weeks (P21 to P42) by which point there is a robust response, followed by doxycycline withdrawal for three weeks (until P63) to allow sufficient time for YAP levels to return to normal (scheme Fig5D). Immunofluorescence against total YAP protein confirmed restoration of the normal YAP expression pattern and levels after recovery (Fig5E), and mRNA *in situ* hybridisation detected a reduction in expression of YAP/TAZ targets *Cyr61* and *Ctgf* (SFig6D). Following recovery from high levels of YAP, the number of SOX2+ cells reduced to comparable levels as in controls (around 10% of the total anterior pituitary) (Fig5E,F). This suggests that the effects of YAP overexpression on the stem cell population are transient following three weeks of treatment (Fig5F).

**Figure 6.**
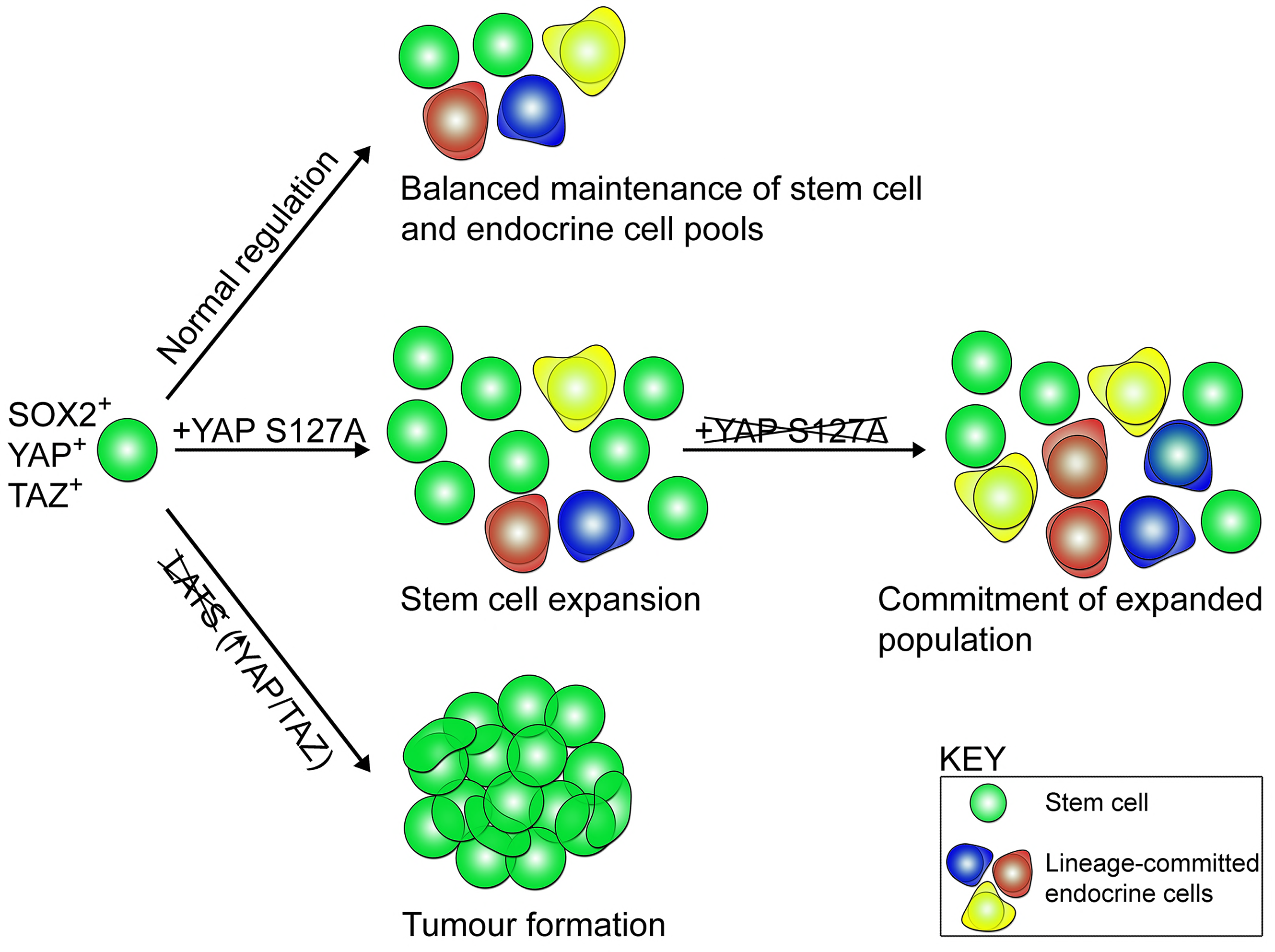
Model of stem cell activity following regulation by the LATS/YAP/TAZ cascade in the anterior pituitary. SOX2+ pituitary stem cells express YAP and TAZ (green spheres). During normal developmental and postnatal expansion (normal regulation), pituitary stem cells are maintained as a balanced pool while generating endocrine cells of three committed lineages (red, blue, yellow). Expression of constitutively active YAP-S127A in pituitary stem cells leads to an expansion of pituitary stem cell numbers and maintenance of the SOX+ state preventing lineage commitment. When YAP-S127A expression ceases, commitment into the endocrine lineages takes place. Genetic deletion of LATS kinases (LATS1 as well as one or two copies of LATS2), results in YAP and TAZ accumulation, repression of lineage commitment, continued expansion of SOX2+ cells and tumour formation.

Finally, to determine if SOX2+ cells could differentiate into hormone-producing cells after the reduction in YAP levels, we expressed constitutive active YAP only in SOX2+ cells whilst lineage tracing this population (*Sox2^CreERT2/+^;R26^rtTA/mTmG^;Col1a1^tetO-Yap^1^/+^*). We induced SOX2+ cells by low-dose tamoxifen administration at P21 and treated with doxycycline for three weeks, followed by doxycycline withdrawal for a further three weeks (Fig5G). Larger clones of SOX2 derivatives were observed at P63 in *Sox2^CreERT2/+^;R26^rtTA/mTmG^;Col1a1^tetO-Yap^1^/+^* animals compared to controls, and these still contained SOX2+ cells (Fig5H). Following withdrawal, we were able to detect GFP+ derivatives of SOX2+ cells, which had differentiated into the three lineages (PIT1, SF1 and ACTH, marking corticotrophs of the TPIT lineage) (Fig5I). Taken together, these findings confirm that sustained expression of YAP is sufficient to maintain the SOX2+ state and promote activation of normal SOX2+ pituitary stem cells *in vivo*, driving expansion of this population.

## DISCUSSION

Here we establish that regulation of LATS/YAP/TAZ signaling is essential during anterior pituitary development and can influence the activity of the stem/progenitor cell pool. LATS kinases, mediated by YAP and TAZ, are responsible for controlling organ growth, promoting an undifferentiated state and repressing lineage commitment. Loss of both *Lats1* and *Lats2*, encoding potent tumour suppressors, leads to dramatic tissue overgrowth during gestation, revealing a function for these enzymes in restricting growth during pituitary development. The involvement of YAP/TAZ and dysfunction of the kinase cascade is emerging in multiple paediatric cancers, which are often developmental disorders ^38^.

Loss of *LATS1* heterozygosity has been reported in a range of human tumours ^39, 40, 41, 42^ leading to an increase in YAP/TAZ protein levels. Previous global deletion of *Lats1* in mice resulted in a variety of soft tissue sarcomas and stromal cell tumours ^43^. The anterior lobe of these animals appeared hyperplastic with poor endocrine cell differentiation leading to combined hormone deficiencies, but the presence of tumours was not noted. We report that loss of *Lats1* alone is sufficient to drive anterior and intermediate lobe tumour formation. This phenotype is accelerated following additional deletion of one copy of *Lats2*. Phenotypically identical tumour lesions were generated when the genetic deletions were carried out embryonically in the RP, or at postnatal stages. Interestingly, tissue-specific loss of *Mst1* and *Mst2*, which regulate LATS activation in other tissues ^44^, did not lead to any pituitary defects. Similar situations have been reported in other organs where LATS are functioning ^44^. These data suggest that alternative regulation on LATS kinases is likely to be active in the pituitary. The resulting non-secreting tumours in our mouse models are composed predominantly of SOX2+ stem cells and display signs of squamous differentiation. Rare cases of squamous cell carcinoma have been reported as primary pituitary tumours ^45^, but more frequently, arising within cysts that are normally non-neoplastic epithelial malformations ^46, 47^. Although human pituitary carcinomas are only diagnosed as such after metastasis, the tumours generated in our mouse models fit their histopathological profile. Genetic lineage tracing identified SOX2+ cells as the cell of origin of the tumours; this observation could have ramifications regarding involvement of the LATS/YAP/TAZ pathway in the establishment or progression of human pituitary tumours composed of uncommitted cells. In cancer stem cells of osteosarcoma and glioblastoma, SOX2 antagonises upstream Hippo activators, leading to enhanced YAP function^48^. We recently reported enhanced expression of YAP/TAZ in a range of non-functioning human pituitary tumours, compared to functioning adenomas, and that *Lats1* knock-down in GH3 pituitary mammosomatotropinoma cells results in repression of the *Gh* and *Prl* promoters ^29^. Therefore, YAP/TAZ, perhaps in a positive feedback loop with SOX2, are likely to function both to promote the maintenance of an active pituitary stem cell state as well as to inhibit differentiation.

By dissecting the downstream requirement for YAP in pituitary regulation by the LATS/YAP/TAZ axis, we found that expression of constitutively active YAP (S127A) is sufficient to push SOX2+ pituitary stem cells into an activated state, leading to expansion of the stem cell cohort. YAP has previously been indicated to promote the stem cell state in other tissues, e.g. pancreas neurons mammary glands ^49^. However, this does not fully recapitulate the LATS deletion phenotypes, as it did not lead to the formation of tumours during the time course of YAP activation (12 weeks). However, the temporal control of expressing the mutation is critical, as seen in other tumour models ^50^. Instead, the findings identify an isolated role for YAP in promoting the expansion of the SOX2+ stem cell pool and restoring their proliferative potential to levels akin to the most active state during postnatal pituitary growth. Activity of YAP/TAZ is reduced in dense tissues, resulting in a decrease in stemness. One mechanism through which this is achieved is by crosstalk with other signaling pathways regulating stem cell fate^51, 52^. For example, a decrease in YAP/TAZ activity removes inhibition on Notch signalling, resulting in higher levels of differentiation and a drop in stem cell potential ^53^. In the pituitary, Notch plays a role in the maintenance of the SOX2 stem cell compartment and is involved in regulating differentiation ^54, 55, 56, 57^. The downstream mechanisms of YAP action on SOX2+ pituitary stem cells, as well as the likely crosstalk with other signalling pathways remain to be explored.

In summary, our findings highlight roles for LATS/YAP/TAZ in the regulation of pituitary stem cells, where fine-tuning of their expression can make the difference between physiological stem cell re-activation and tumourigenesis, of relevance to other organs. We reveal this axis is involved in the control of cell fate commitment, regulation of regenerative potential and promotion of tumourigenesis. These findings can aid in the design of treatments against pituitary tumours and in regenerative medicine approaches targeting the regulation of endogenous stem cells.

## AUTHOR CONTRIBUTIONS

Conceptualization C.L.A.; Methodology E.J.L., C.L.A.; Investigation E.J.L., J.P.R., A.S, P.X., T.S.J., S.T.; Resources R.L.J.; Writing – Original Draft, C.L.A.; Writing-Review & Editing C.L.A. and E.J.L.; Supervision C.L.A.; Funding Acquisition, C.L.A. and S.R.B.

### ACKNOWLEDGEMENTS

This study has been supported by grant MR/L016729/1 from the MRC and a Lister Institute Research Prize to C.L.A., by the Deutsche Forschungsgemeinschaft (DFG) within the CRC/Transregio 205/1 as well as GRK 2251 to C.L.A. and S.R.B. E.J.L. was supported by the King’s Bioscience Institute and the Guy’s and St Thomas’ Charity Prize PhD Programme in Biomedical and Translational Science. J.P.R. was supported by a Dianna Trebble Endowment Fund Dental Institute Studentship. We thank Prof. Jacques Drouin and Prof. Simon Rhodes for TPIT and PIT1 antibodies respectively, and the National Hormone and Peptide Program (Harbor–University of California, Los Angeles Medical Center) for providing some of the hormone antibodies used in this study. We thank Prof. Juan Pedro Martinez-Barbera, Dr Rocio Sancho and Dr Marika Charalambous for discussions and critical reading of the manuscript. The authors declare no conflict of interest.

## MATERIALS & METHODS

### Animals

Animal husbandry was carried out under compliance of the Animals (Scientific Procedures) Act 1986, Home Office license and KCL ethical review approval.

The *Hesx1^Cre/+^* ^58^, *Sox2^CreERT2^/+* ^1^, *Yap*^*fl/fl*^ ^26^, *Taz^-/-^* ^31^ (JAX:011120), *R26^mTmG/+^* ^59^(JAX:007576), *ROSA26^rtTA/+^* ^60^ (JAX:016999), *Col1a1^tetO-Yap1^/+* ^61^(MGI:5430522), *Mst1^fl/fl^;Mst2^fl/fl^* ^62^(JAX:017635), and *Lats1^fl/fl^* ^52^(JAX:024941) and *Lats2^fl/fl^* ^52^(JAX:025428) have been previously described.

Tamoxifen (Sigma, T5648) was administered to experimental mice by intraperitoneal injection at a single dose of 0.15mg/g body weight, or two equal doses on sequential days, depending on the experiment. Mice for growth studies were weighed every week. For embryonic studies, timed matings were set up where noon of the day of vaginal plug was designated as 0.5dpc.

For YAP-TetO experiments, crosses between *Hesx1^Cre/+^;R26^+/+^;Col1a1^+/+^* and *Hesx1^+/+^;R26^rtTA/rtTA^;Col1a1^tetO-Yap1^/ tetO-Yap^1^* animals were set up to generate *Hesx1^Cre/+^;R26^rtTA/+^;Col1a1^tetO-Yap^1^/+^* offspring (hereby YAP-TetO) and control littermates, or crosses between *Sox2^CreERT^2^/+^;R26^mTmG/mTmG^;Col1a1^+/+^* and *Sox2^+/+^; R26^rtTA/rtTA^;Col1a1^tetO-Yap^1^/ tetO-Yap1^* animals were set up to generate *Sox2^CreERT^2^/+^;R26^rtTA/mTmG^;Col1a1^tetO-Yap^1^/+^* offspring.

Whilst treated with the tetracycline analogue doxycycline, YAP-TetO expressed rtTA from the *ROSA26* locus in *Cre*-derived cells, enabling YAP S127A expression from the *Col1a1* locus. For embryonic studies between 5.5dpc and 15.5dpc (scheme, Fig1A), doxycycline (Alfa Aesar, J60579) was administered to pregnant dams in the drinking water at 2mg/ml, supplemented with 10% sucrose. For postnatal analyses animals were treated with doxycycline or vehicle (DMSO) as described, from the ages specified for individual experiments on the *Hesx1^Cre/+^* driver, or directly following tamoxifen administration for animals on the *Sox2^CreERT^2^/+^* driver.

### Tissue preparation

Embryos and adult pituitaries were fixed in 10% neutral buffered formalin (Sigma) overnight at room temperature. The next day, tissue was washed then dehydrated through graded ethanol series and paraffin-embedded. Embryos up to 13.5dpc were sectioned sagittal and all older embryo and postnatal samples were sectioned frontal, at a thickness of 7µm for immunofluorescence staining, or 4µm for RNAscope mRNA *in situ* hybridisation.

### **RNAscope mRNA *in situ* hybridisation**

Sections were selected for the appropriate axial level, to include Rathke’s pouch or pituitary, as described previously ^28^. The RNAscope 2.5 HD Reagent Kit-RED assay (Advanced Cell Diagnostics) was used with specific probes: *Ctgf, Cyr61, Lats2* (all ACDBio).

### H&E staining

Sections were dewaxed in histoclear and rehydrated through graded ethanol series from 100% to 25% ethanol, then washed in distilled H_2_O. Sections were stained with Haematoxylin QS (Vector #H3404) for 1 minute, and then washed in water. Slides were then stained in eosin in 70% ethanol for 2 minutes and washed in water. Slides were dried and coverslips were mounted with VectaMount permanent mounting medium (Vector Laboratories H5000).

### Immunofluorescence and immunohistochemistry

Slides were deparaffinised in histoclear and rehydrated through a descending graded ethanol series. Antigen retrieval was performed in citrate retrieval buffer pH6.0, using a Decloaking Chamber NXGEN (Menarini Diagnostics) at 110°C for 3mins. Tyramide Signal Amplification (TSA) was used for staining using antibodies against YAP (1:1000, Cell Signaling #4912S), pYAP (1:1000, Cell Signaling #4911S), TAZ (1:1000, Atlas Antibodies #HPA007415) and SOX2 (1:2000, Abcam ab97959) with EMCN (1:1000, Abcam ab106100) staining as follows: sections were blocked in TNB (0.1M Tris-HCl, pH7.5, 0.15M NaCl, 0.5% Blocking Reagent (Perkin Elmer FP1020)) for 1 hour at room temperature, followed by incubation with primary antibody at 4C overnight, made up in TNB. Slides were washed three times in TNT (0.1MTris-HCl pH7.5, 0.15M NaCl, 0.05% Tween-20) then incubated with secondary antibodies (biotinylated anti-rabbit (1:350 Abcam ab6720) and anti-Rat Alexa Fluor 555 (1:300, Life Technologies A21434) for 1 hour at room temperature and Hoechst (1:10000, Life Technologies H3570). Slides were washed again then incubated in ABC reagent (ABC kit, Vector Laboratories PK-6100) for 30 mins, followed by incubation with TSA conjugated fluorophore (Perkin Elmer NEL753001KT) for ten minutes. Slides were washed and mounted with VectaMount (Vector Laboratories H1000).

For regular immunofluorescence sections were blocked in blocking buffer (0.15% glycine, 2mg/ml BSA, 0.1% Triton-X in PBS), with 10% sheep serum (donkey serum for goat SOX2 antibody) for 1 hour at room temperature, followed by incubation with primary antibody at 4C overnight, made up in blocking buffer with 1% serum. Primary antibodies used were against SOX2 (1:250, Immune Systems Ltd GT15098), active YAP (1:300, Abcam ab205270), GFP (1:300, Abcam ab13970), Ki-67 (1:300, Abcam ab16667), SOX9 (1:300, Abcam ab185230), PIT1 (1:1000, Gift from S. Rhodes, Indiana University), TPIT (1:1000, Gift from J. Drouin, Montreal), SF1 (1:200, Life Technologies N1665), Gamma H2A.X (1:1000, Abcam ab2893), Vimentin (1:300, Cell Signaling #5741), Caspase (1:300, Cell Signaling #9661S). Slides were washed in PBST then incubated with secondary antibodies for 1 hour at room temperature. Appropriate secondary antibodies were incubated in blocking buffer for 1 hr at room temperature (biotinylated anti-rabbit (1:350, Abcam ab6720), biotinylated anti-mouse (1:350, Abcam ab6788), anti-chicken 488 (1:300, Life Technologies A11039), anti-goat 488 (1:300, Abcam ab150133). Slides were washed again using PBST and incubated with fluorophore-conjugated Streptavidin (1:500, Life Technologies S21381 or S11223) for 1 hour at room temperature, together with Hoechst (1:10000, Life Technologies H3570). Slides were washed in PBST and mounted with VectaMount (Vector Laboratories, H1000).

Immunohistochemistry for the remaining antigens were undertaken on a Ventana Benchmark Autostainer (Ventana Medical Systems) using the following primary antibodies and antigen retrieval: AE1/AE3 (1:100, Dako M351529), CC1 (36 minutes, Ventana Medical Systems 950-124); Chromogranin (1:400, Dako M086901), CC1 (36 minutes, Ventana Medical Systems 950-124); NCAM (1:15, Novocastra NCL-L-CD56-504), CC1 (64 minutes, Ventana Medical Systems 950-124); NSE (1:1000, Dako M087329), CC1 (36 minutes, Ventana Medical Systems 950-124); p63 (1:100, A. Menarini Diagnostics), CC1 (64 minutes, Ventana Medical Systems 950-124) and Synaptophysin (1:2, Dako M731529), CC2 (92 minutes, Ventana Medical Systems 950-124). Targets were detected and viewed using the ultraView Universal DAB Detection Kit (Ventana Medical Systems, 760-500) according to manufacturer’s instructions.

### Imaging

Wholemount images were taken with a MZ10 F Stereomicroscope (Leica Microsystems), using a DFC3000 G camera (Leica Microsystems). For bright field images, stained slides were scanned with Nanozoomer-XR Digital slide scanner (Hamamatsu) and images processed using Nanozoomer Digital Pathology View. Fluorescent staining was imaged with a TCS SP5 confocal microscope (Leica Microsystems) and images processed using Fiji ^63^.

### Quantifications of cell number

Cell counts were performed manually using Fiji cell counter plug-in; 5-10 fields were counted per sample, totalling over 1500 nuclei, across 3-5 pituitaries. Statistical analyses and graphs were generated in GraphPad Prism (GraphPad Software).

## SUPPLEMENTARY FIGURE LEGENDS

**Supplementary Figure 1 Regulation of YAP and TAZ during pituitary development.**

**A.** Hematoxylin and eosin staining on frontal sections through the pituitary from control and *YAP-TetO* heads after DOX treatment from 5.5dpc until 15.5dpc. **B.** Hematoxylin and eosin staining on sagittal pituitary sections of 13.5dpc *Hesx1^Cre/+^;Yap^fl/fl^;Taz^-/-^* (mutant) and *Hesx1^+/+^;Yap^fl/+^;Taz^+/-^* (control) showing comparable morphology. **C.** Immunofluorescence staining using antibodies against SOX2 in *Hesx1^Cre/+^;Yap^fl/fl^;Taz^-/-^* and control at 13.5dpc (sagittal) and 16.5dpc (frontal) showing the presence of SOX2+ cells in both genotypes. **D.** Immunofluorescence staining for SOX2, endomucin (EMCN) and active YAP in P28 *Hesx1^Cre/+^;Yap^fl/fl^;Taz^-/-^* and control pituitaries, identifies SOX2+ cells in regions that are negative for active YAP (mice are null for TAZ) and normal vasculature. **E.** Immunofluorescence staining for lineage committed progenitor markers PIT1 and SF1, as well as ACTH marking corticotrophs (TPIT lineage), reveals the presence and normal localisation of cells from the three lineages in a P28 *Hesx1^Cre/+^;Yap^fl/fl^;Taz^-/-^* mutant. Scale bars 1mm in A, 100µm in B-E.

**Supplementary Figure 2 Pituitary-specific loss of *Mst1* and *Mst2* does not affect SOX2 cell specification or lineage commitment.**

**A.** Dorsal view of wholemount *Hesx1^Cre/+^;Mst1^fl/fl^;Mst2^fl/fl^* (mutant) and *Mst1^fl/fl^;Mst2^fl/fl^* (control) pituitaries at P0 showing comparable morphology and size at birth. **B.** Immunofluorescence staining using antibodies against SOX2, TAZ, endomucin (EMCN), YAP and pYAP at P0, indicating comparable staining between control and mutant samples. **C.** Immunofluorescence staining against lineage commitment markers PIT1, TPIT and SF1 shows normal lineage commitment in a *Hesx1^Cre/+^;Mst1^fl/fl^;Mst2^fl/fl^* mutant pituitary compared to the control at P10. **D.** Hematoxylin and eosin staining through frontal sections of *Hesx1^Cre/+^;Mst1^fl/fl^;Mst2^fl/fl^* and control pituitaries at P70. AL: anterior lobe, IL: intermediate lobe, PL: posterior lobe. Scale bars 100µm.

**Supplementary Figure 3 Isolated deletions of *Lats1* or *Lats2* in the pituitary do not affect development.**

**A.** Hematoxylin and eosin staining of a sagittal section of *Hesx1^Cre/+^;Lats1^fl/fl^* at 13.5dpc showing normal morphology (see Figure 2A for control). Dashed lines demarcate developing Rathke’s pouch. Immunofluorescence staining for TAZ and YAP reveals a normal expression pattern and no gross protein accumulation (compare to control, Figure 2A) **B.** Dorsal view of wholemount *Hesx1^Cre/+^;Lats1^fl/fl^* (mutant) and *Hesx1^Cre/+^* (control) pituitaries at P0 showing comparable morphology and size at birth. **C.** RNAscope mRNA *in situ* hybridisation against *Lats2* shows an increase in transcripts in the anterior pituitary following deletion of *Lats1* (*Hesx1^Cre/+^;Lats1^fl/fl^*) compared to control (*Hesx1^Cre/+^*), where *Lats2* expression is barely detectable**. D.** Hematoxylin and eosin staining of a sagittal section of *Hesx1^Cre/+^;Lats2^fl/fl^* at 13.5dpc showing normal morphology (see Figure 2A for control). Dashed lines demarcate developing Rathke’s pouch. **E.** Hematoxylin and eosin staining on frontal sections through 15.5dpc embryonic heads of *Hesx1^Cre/+^;Lats1^fl/fl^;Lats2^fl/fl^* (mutant) and control (*Hesx1^+/+^;Lats1^fl/fl^;Lats2^fl/fl^*) genotypes, at the levels indicated in the cartoon. Note the hyperplastic pituitary at both axial levels, exerting mass effect on the brain. Asterisk indicates necrosis. Scale bars 100µm in A-D, 1mm in E.

**Supplementary Figure 4 Analysis of tumourigenic lesions in postnatal pituitaries following pituitary-specific deletion of *Lats1.***

**A.** Immunofluorescence staining for TAZ and active YAP reveal lesions of accumulation at P21 in *Hesx1^Cre/+^;Lats1^fl/fl^;Lats2^fl/+^*compared to *Hesx1^+/+^;Lats1^fl/fl^;Lats2^fl/+^*control. Immunofluorescence staining using antibodies against SOX2 and endomucin (EMCN) show these lesions are composed of SOX+ stem cells and have reduced vascularisation. **B.** Hematoxylin and eosin staining of frontal sections from *Hesx1^Cre/+^;Lats2^fl/fl^* and *Hesx1^Cre/+^* control pituitaries at P56 showing comparable histology. **C.** Immunofluorescence staining against SOX2 and endomucin on an intermediate lobe lesion (asterisk) in a *Hesx1^Cre/+^;Lats1^fl/fl^;Lats2^fl/+^* pituitary compared to control. **D.** Immunofluorescence staining against DNA damage marker gamma H2A.X showing positive cells in *Hesx1^Cre/+^;Lats1^fl/fl^;Lats2^fl/+^*mutants. **E.** P56 Immunohistochemistry using antibodies against p63 and the AE1/AE3 cytokeratin cocktail, both markers of pituitary carcinomas, showing abundant staining in *Hesx1^Cre/+^;Lats1^fl/fl^;Lats2^fl/+^* compared to control. **F.** Immunohistochemistry using antibodies against synaptophysin, neural-specific enolase (NSE) and chromogranin demonstrate tumourigenic lesions in *Hesx1^Cre/+^;Lats1^fl/fl^;Lats2^fl/+^* are negative for adenoma markers. Lesions are negative for vimentin by immunofluorescence staining, commonly marking spindle-cell oncocytoma in the pituitary. Scale bars 100µm in A, C-F; 500µm in B. PL: posterior lobe, IL: intermediate lobe, AL: anterior lobe.

**Supplementary Figure 5 Analysis of tumourigenic lesions in postnatal pituitaries following SOX2-specific deletion of *Lats1.***

**A.** Immunohistochemistry using specific antibodies against p63 and cytokeratin cocktail AE1/AE3 on frontal sections of *Sox2^CreERT^2^/+^;Lats1^fl/fl^;Lats2^fl/+^* (mutant) and *Sox2^+/+^;Lats1^fl/fl^;Lats2^fl/+^* (control) pituitaries at P35, revealing positive staining in mutants. **B.** Double immunofluorescence staining against total YAP and GFP, as well as SOX2 and GFP in consecutive sections of a tumourigenic lesion from *Sox2^CreERT^2^/+^;Lats1^fl/fl^;Lats2^fl/+^*;R26^*mTmG/+*^ pituitaries at P35. Lineage tracing of SOX2+ cells, detected using GFP reveals abundant staining in the tumour lesion, characterised by accumulation of YAP and SOX2+ cells (yellow arrowheads). Scale bars 100µm.

**Supplementary Figure 6 Postnatal expression of constitutively active YAP increases leads to an activation of SOX2+ pituitary stem cells.**

**A.** Schematic outlining the time course of doxycycline (DOX) treatment administered to *Hesx1^Cre/+^;R26^rtTA/+^;Col1a1^tetO-Yap^1^/+^* (*YAP-TetO*) and *Hesx1^+/+^;R26^rtTA/+^;Col1a1^tetO-Yap^1^/+^* controls to drive expression of YAP-S127A in mutant pituitaries. Hematoxylin and eosin staining of control and *YAP-TetO* pituitaries at P42 (3 weeks treatment), P63 (6 weeks treatment) and P105 (12 weeks treatment). **B.** RNAscope mRNA *in situ* hybridisation against YAP targets *Cyr61* and *Ctgf* showing increased transcripts in *YAP-TetO* sections compared to controls at P42. **C.** Analysis of *YAP-TetO* mutants at P105: double immunofluorescence staining against SOX2 and Ki-67 reveals regions of expanded SOX2+;Ki-67-cells compared to the normal expression pattern in the control. This region is SOX9+, does not accumulate TAZ or YAP and expresses pYAP as does normal anterior pituitary epithelium. Immunofluorescence against PIT1 shows the absence of commitment to this lineage, a pattern not seen in the control. Hematoxylin and eosin staining in consecutive sections identifies this region, which does not have neoplastic features. **D.** Schematic outlining the time course of doxycycline (DOX) treatment administered to *Hesx1^Cre/+^;R26^rtTA/+^;Col1a1^tetO-Yap^1^/+^* (*YAP-TetO*) and *Hesx1^+/+^;R26^rtTA/+^;Col1a1^tetO-Yap^1^/+^* controls to drive expression of YAP-S127A in mutant pituitaries for three weeks, followed by a three-week recovery period in the absence of DOX. Hematoxylin and eosin staining of control and *YAP-TetO* pituitaries. RNAscope mRNA *in situ* hybridisation shows comparable levels of expression of targets *Cyr61* and *Ctgf.* Scale bars 250µm in A, 100µm in B-D.

